# Systemic administration of PD-L1 blocking antibodies leads to removal of senescent microglia

**DOI:** 10.64898/2025.12.22.695923

**Authors:** Yuliya Androsova, Alexander Kertser, Hannah Partney, Bar Nathansohn, Angham Ibraheem, Miguel Abellanas, Tommaso Croese, Tomer Meir Salame, Hagay Akiva, Valery Krizhanovsky, Michal Schwartz

## Abstract

Senescent microglia develop during aging and Alzheimer’s disease (AD), driving chronic neuroinflammation. Here we hypothesized that the previously observed disease-modifying effects of PD-1/PD-L1 blockade occur through clearance of senescent microglia. Using CyTOF, we found that a single systemic anti-PD-L1 injection leads to rapid elimination of senescent microglia in 5xFAD and aged wild-type mice, independently of Fc effector function, while increasing homeostatic microglia. These findings suggest that immune rejuvenation via PD-L1 blockade promotes disease modification in AD through senescent-microglial elimination.

## Introduction

Brain function and repair are tightly dependent on support from the peripheral immune system^1^. Thus, aging of the immune system contributes to cognitive decline and impaired brain function^2–4^. Aging is also recognized as a major risk factor for late-onset dementia, and particularly Alzheimer’s disease (AD). Though the primary cause of AD and other neurodegenerative diseases is often linked to accumulation of misfolded proteins, local brain inflammation acts as a catalyst of symptom onset and disease progression^5–7^. Accordingly, therapies targeting misfolded protein accumulation, such as amyloid plaques, have proven insufficient for meaningful disease modification, even if biologically effective^8^.

The neuroinflammation that emerges during disease progression is associated, at least in part, with microglial dysfunction. While microglia can normally contain perturbations arising from deviations in brain homeostasis, persistent exposure to the threats may drive accelerated proliferation of microglia^9^, followed by loss of function, exhaustion and acquisition of a senescent state^10^. Senescent cells are characterized by stable cell-cycle arrest, persistent DNA damage response, resistance to apoptosis, and expression of Senescence-Associated-Secretory-Phenotype (SASP)^11^. This pro-inflammatory activity drives chronic tissue inflammation through continuous secretion of pro-inflammatory cytokines^12^. Notably, senescent microglia demonstrate a conserved molecular signature across various brain conditions, including aging and amyloidosis^13–15^.

Targeting the PD-1/PD-L1 inhibitory checkpoint pathway has been shown to confer disease-modifying benefits across multiple models of neurodegeneration, including AD and tauopathy^16–19^. Importantly, one of the earliest detectable effects following a single anti-PD-L1 administration in animal models of dementia is a reduction in local brain inflammation^16–20^. We therefore hypothesized that this early anti-inflammatory response results from clearance of senescent microglia, with additional therapeutic benefits occurring downstream to this primary activity.

Here, we show that in the 5xFAD mouse model of AD, the same immunotherapy that was previously shown to modify disease progression, induces clearance of senescent microglia, detectable as early as 3 days following a single systemic anti-PD-L1 injection.

## Results

### A single injection of anti-PD-L1 antibody leads to a reduction in senescent microglia in both 5xFAD and aged mice

We first assessed the effect of checkpoint blockade on senescent microglia, using 7-8-month-old and 10-11-month-old 5xFAD mice (Fig. 1a). Antibody dosing was selected based on our previous studies^16,19^. In the initial experiment, 5xFAD mice received a single intraperitoneal injection of 1.5 mg anti-PD-L1, or a control antibody against an irrelevant epitope (anti-KLH). Mice were euthanized 12 days post-injection, corresponding to the earliest time point previously analysed following a single anti-PD-L1 administration, at which reduction in neuroinflammation was observed^16,17,21^. Brains were excised and processed for cytometry by time-of-flight (CyTOF) analysis^13^. Marker panels were selected based on previous studies, enabling discrimination between homeostatic, activated and senescent microglial states (Table 1).

**Figure 1.**
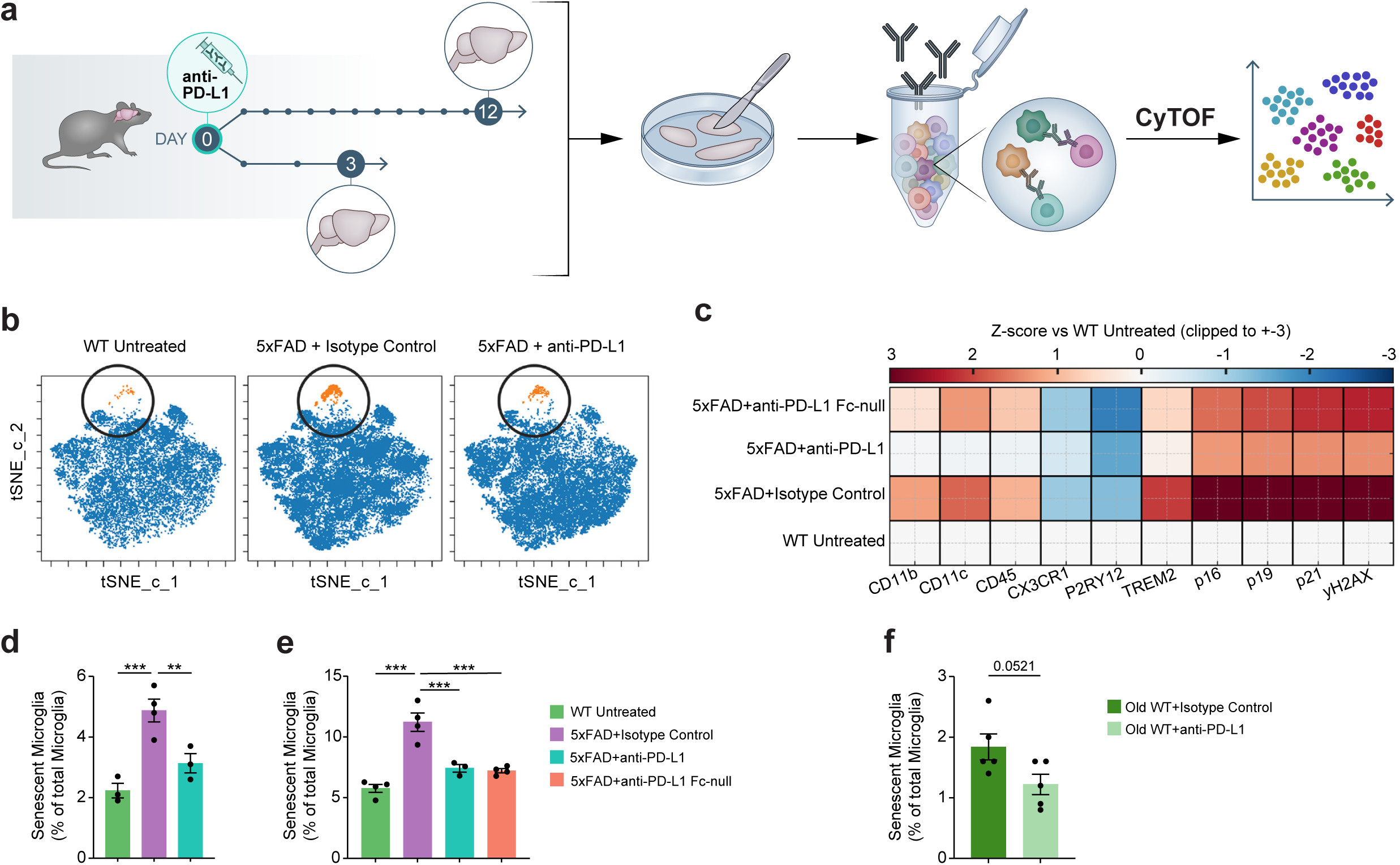
Dynamics of senescent microglia in 5xFAD mice following anti-PD-L1 treatment. **a.** Experimental design. Cohorts of 5xFAD and C57BL/6 (wild-type, WT) mice received either a single intraperitoneal injection of anti-PD-L1 (1.5 mg), an IgG2 isotype control (1.5 mg), or an Fc-null surrogate anti-PD-L1 antibody. Brains were collected at day 3 or day 12 post-injection, enzymatically digested, stained, and analysed by CyTOF. **b.** Representative t-SNE projections of microglia (4,099 randomly sampled cells per sample), identified as CD45⁺CD11b⁺CX3CR1⁺P2RY12⁺Ly6C/G⁻CD206⁻CCR2⁻. Senescent microglia (orange) were defined as CD45⁺CD11b⁺CX3CR1⁺P2RY12⁺Ly6C/G⁻CD206⁻CCR2⁻Ki67⁻p16⁺TREM2⁺p19⁺p21⁺γH2AX⁺. Clustering was performed using FlowSOM in Cytobank. **c.** Heatmap of median marker expression in the microglial cluster, Z-score normalized to WT untreated controls. **d.** CyTOF quantification of senescent microglia at day 3 post-treatment in 7–8-month-old 5xFAD mice treated with anti-PD-L1 or isotype control, compared to WT. One-way ANOVA, *P = 0.0019*; Fisher’s LSD (uncorrected): WT untreated vs isotype, *P* < 0.001; isotype vs anti-PD-L1, *P* = 0.008. **e.** CyTOF quantification of senescent microglia at day 12 post-treatment in 10-11-month-old 5xFAD mice treated with anti-PD-L1, Fc-null surrogate anti-PD-L1, or isotype control, compared to WT. One-way ANOVA, *P* < 0.001; Brown–Forsythe, *P* = 0.04; Bartlett’s, *P* = 0.007. Fisher’s LSD (uncorrected): WT untreated vs isotype, *P* < 0.001; isotype vs anti-PD-L1, *P* < 0.001; isotype vs Fc-null, *P* < 0.001. **f.** Flow cytometry quantification of senescent microglia (defined as defined as CD45⁺CD11b⁺Ly6C⁻Ki67⁻Bcl-xL⁺γH2AX⁺) in aged WT mice (18–19 months old) at day 12 post-treatment. Unpaired two-tailed t-test, *P* = 0.052.

**Table 1:** Assignment of microglial states based on expression of multiple markers.

|  |  |
| --- | --- |
| Homeostatic microglia | CD45<sup>low</sup>CD11b<sup>+</sup>; CX3CR1<sup>+</sup>; P2RY12<sup>+</sup>; Ly6C/G<sup>-</sup>; CD206<sup>-</sup>; CCR2<sup>-</sup>; Ki67<sup>-</sup>; p16<sup>-</sup>; TREM2<sup>low</sup>; p19<sup>-</sup>; p21<sup>-</sup>; γH2AX<sup>-</sup> |
| Activated microglia | CD45<sup>+</sup>CD11b<sup>+</sup>; CX3CR1<sup>low</sup>; P2RY12<sup>low</sup>; Ly6C/G<sup>-</sup>; CD206<sup>-</sup>; CCR2<sup>-</sup>; CD11c<sup>+</sup>; p16<sup>-</sup>; p19<sup>-</sup>; p21<sup>-</sup>; γH2AX<sup>-</sup> |
| Senescent microglia | CD45<sup>+</sup>CD11b<sup>+</sup>; CX3CR1<sup>+</sup>; P2RY12<sup>+</sup>; Ly6C/G<sup>-</sup>; CD206<sup>-</sup>; CCR2<sup>-</sup>; Ki67<sup>-</sup>; p16<sup>+</sup>; TREM2<sup>+</sup>; p19<sup>+</sup>; p21<sup>+</sup>; γH2AX<sup>+</sup> |

Consistent with our previous observations,^13^ we found accumulation of senescent microglia in 5xFAD mice in an age-dependant manner, reaching 5-10% of the total microglial population by age 7–8-month-old (Fig. 1b-e). A single injection of anti–PD-L1 antibody led to profound reduction in the senescent microglia population compared with age-matched 5xFAD mice receiving isotype-control antibody (Fig. 1d). Prior mechanistic studies in preclinical AD models demonstrated that therapeutic efficacy does not require continuous antibody exposure or Fc-dependant activity, including antibody-dependent cellular cytotoxicity (ADCC) or complement dependent cytotoxicity (CDC). Accordingly, we evaluated the effect of an anti-PD-L1 on clearance of senescent cells using an antibody variant engineered with 4-point mutations that shorten half-life and eliminate Fc-mediated effector function (Fc-null)^21^. The Fc-null anti-PD-L1 reduced senescent microglia to an extent comparable to the native anti-PD-L1 antibody (Fig. 1e), indicating that clearance of senescent microglia also occurs independently of Fc-mediated cytotoxic mechanisms.

We next validated the CyTOF findings using conventional flow cytometry (Extended Data Fig. 1a-c). Senescent microglia were identified by Bcl-xL and γH2AX expression, while excluding Ki-67–positive cells to restrict the analysis to non-proliferating cells exhibiting senescence-associated features (Extended Data Fig. 1a). Using this approach, we tested whether anti-PD-L1 similarly affects senescent microglia in the absence of neurodegenerative pathology. A single injection of anti-PD-L1 was associated with a reduction in the population of senescent microglia in 18-month-old wild-type mice (Fig. 1f). Although CyTOF and flow cytometry yield different absolute cell counts due to inherent methodological differences, both platforms consistently demonstrated the same treatment-associated reduction in senescent microglia following anti-PD-L1 administration.

### CCR2 signalling is required for anti-PD-L1 mediated clearance of senescent microglia

In our previous studies, the anti-PD-L1-mediated therapeutic effect was shown to be dependent on the CCR2-CCL2 pathway, and specifically on monocyte-derived macrophages ^16–18^. We therefore assessed here whether the effect on senescent microglia was similarly dependent on this axis. To this end, we blocked CCR2-dependent monocyte egress from the bone marrow using anti-CCR2 antibody administered according to a previously established protocol, starting 3 days before anti-PD-L1 administration, and repeated every 4 days^17^ (Fig. 2a). Under these conditions, the anti-PD-L1-induced reduction in senescent microglia observed 12 days after a single treatment was completely abrogated (Fig. 2b, c).

**Figure 2.**
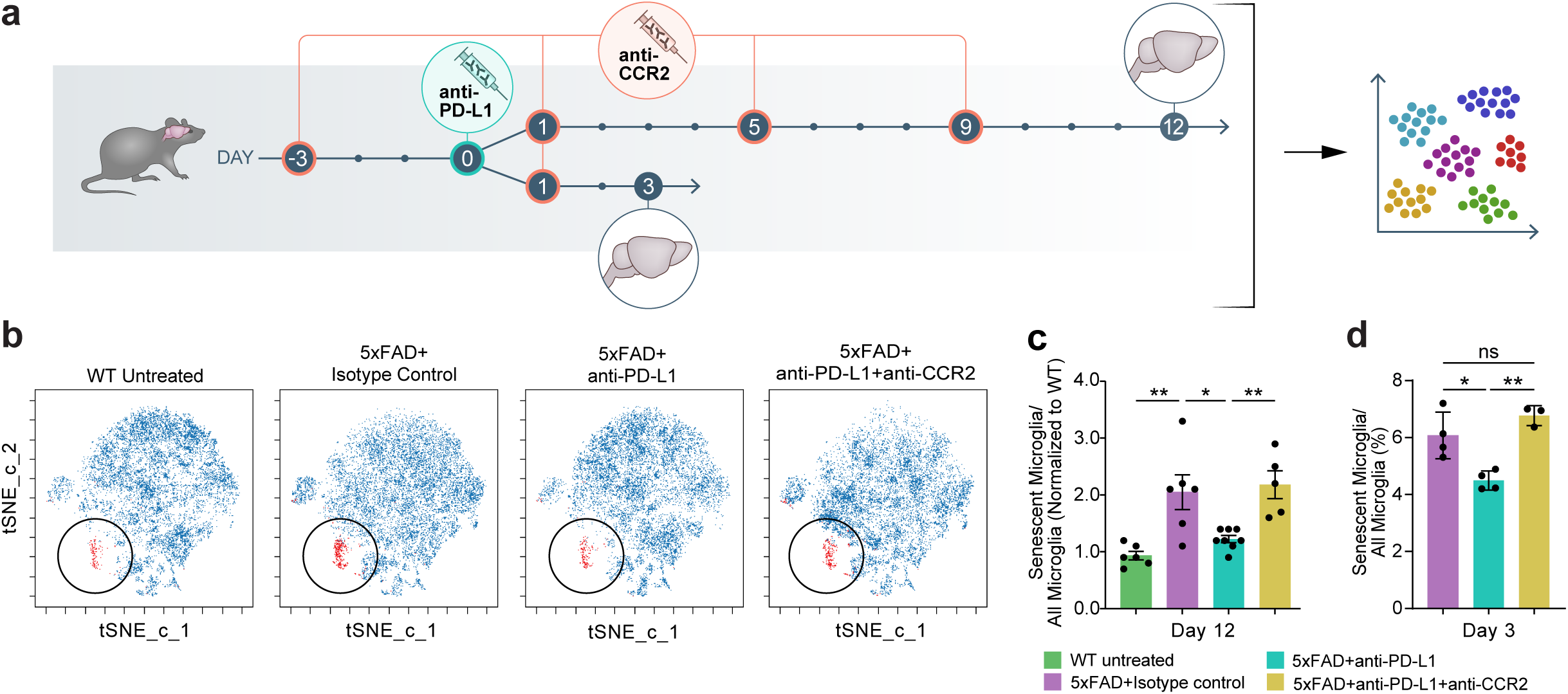
CCR2-dependent clearance of senescent microglia and reshaping of the brain immune landscape by anti-PD-L1. **a.** Experimental design. 5xFAD and WT mice received either (1) a single injection of anti-PD-L1 (1.5 mg), (2) an IgG2 isotype control (1.5 mg), or (3) combination therapy consisting of anti-PD-L1 plus anti-CCR2 (M21, 400 μg per injection, intraperitoneally every 4 days for 4 doses, starting 3 days prior to anti-PD-L1). Brains were collected at day 3 and 12, and analysed by CyTOF. **b.** Representative t-SNE projections of microglia (3,876 randomly sampled cells per sample), identified as in Fig. 1b. Senescent microglia (red) are highlighted within the total microglial population (blue). **c.** CyTOF quantification of senescent microglia at day 12 in 7-8-month-old 5xFAD mice treated with isotype control, anti-PD-L1, or combination anti-PD-L1 + anti-CCR2 in comparison to WT, and in 10–11-month-old mice treated with the same regimens. One-way ANOVA, *P* = 0.0002; Bartlett’s test, *P* = 0.002. Tukey’s multiple comparisons (uncorrected): WT untreated vs isotype (**), isotype vs anti-PD-L1 (*), anti-PD-L1 vs anti-PD-L1+anti-CCR2 (**). Data from two independent experiments were normalized to untreated WT samples, and pooled. **d.** CyTOF quantification of senescent microglia at day 3 in 7–8-month-old 5xFAD mice treated with isotype control, anti-PD-L1, or anti-PD-L1 + anti-CCR2 in comparison to WT. One-way ANOVA, *P = 5.89 × 10⁻⁴* ; Bartlett’s test, *P* = 0.002. Tukey’s multiple comparisons (uncorrected): anti-PD-L1 vs isotype (*), anti-PD-L1 vs anti-PD-L1+anti-CCR2 (**).

Given that CCR2-dependent leukocyte recruitment is an early and transient immunological response, together with the rapid clearance of anti-PD-L1 blocking antibody from the circulation, we hypothesized that senescent microglial clearance represents an earlier event following treatment, rather than a downstream consequence of disease modification. To test this, we repeated the same experiments and excised brains 3 days after anti-PD-L1 administration. Mice were treated with anti-CCR2 three days prior to, and on the day of anti-PDL1 administration (Fig. 2a). Already at this early timepoint, anti-PD-L1 led to a significant reduction of senescent microglia, an effect that was completely abrogated by blocking CCR2 (Fig. 2d). These findings suggest that anti-PD-L1–driven clearance of senescent microglia precedes other disease-modifying effects previously reported in this mouse model^16,18^.

We next assessed whether anti-PD-L1 treatment also alters the incidence of other microglial states beyond senescence. In 5xFAD mice, we quantified the levels of homeostatic and proliferating microglia following anti-PD-L1 administration. At 12 days after treatment, pooled analysis from two independent experiments revealed an increased proportion of homeostatic microglia within the total microglial compartment (Fig. 3a). Consistent with previous reports ^17^, proliferating microglia were elevated in untreated 5xFAD mice, and were markedly reduced by anti-PD-L1 treatment to levels comparable to those in WT mice (Fig. 3b).

**Figure 3.**
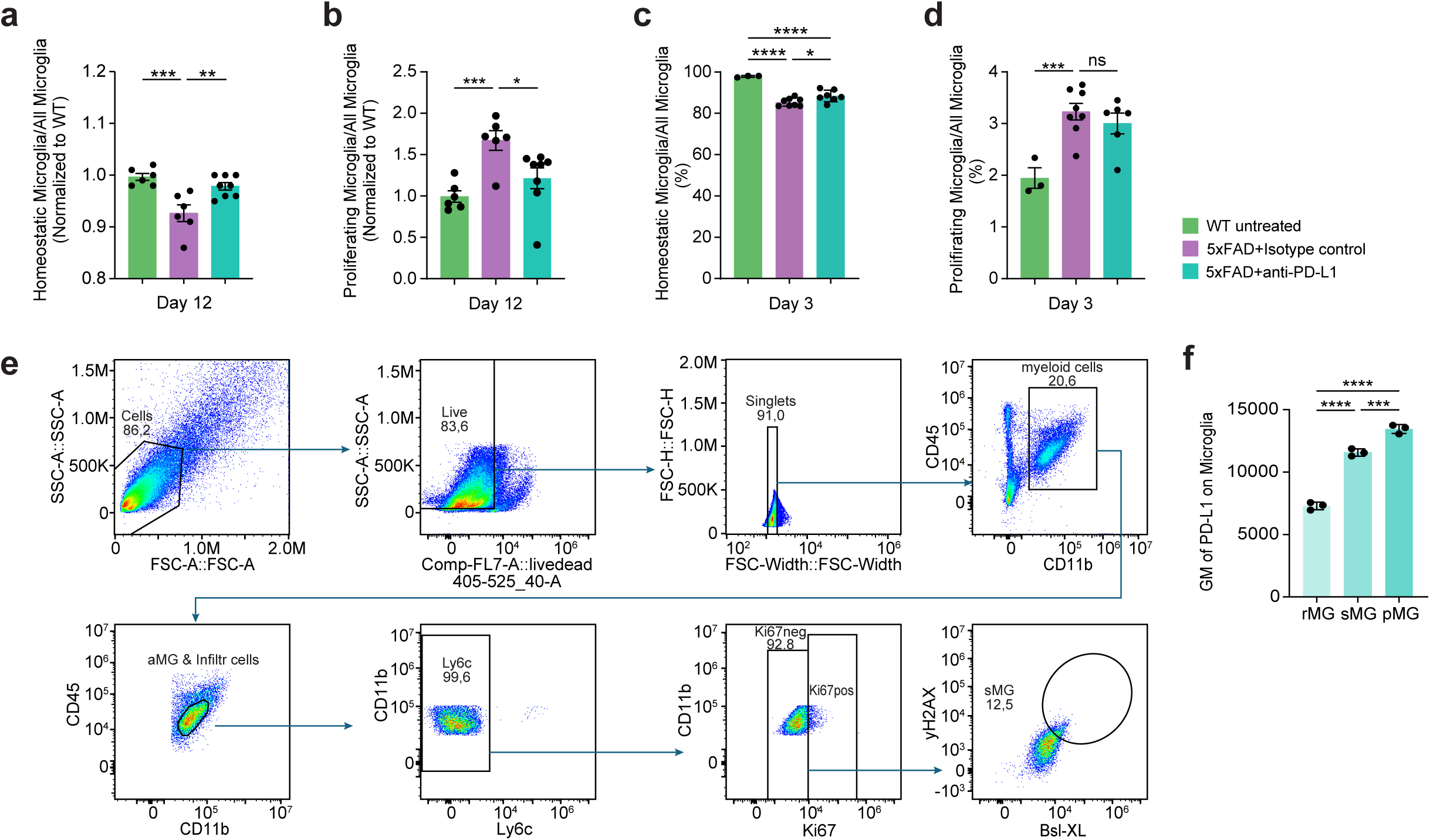
Anti-PD-L1 reshapes the brain - immune landscape. **a.** CyTOF quantification of homeostatic microglia (CD45⁺CD11b⁺CX3CR1^hiP2RY12^hiLy6C/G⁻CD206⁻CCR2⁻Ki67⁻p16⁻TREM2⁻p19⁻p21⁻γH2 AX⁻) at day 12 in 7-8-month-old 5xFAD mice treated with isotype control or anti-PD-L1, compared to WT, and in 10–11-month-old mice treated with the same regimens. One-way ANOVA, *P* < 0.001; Fisher’s LSD (uncorrected): WT vs isotype, *P* < 0.001; isotype vs anti-PD-L1, *P* = 0.002. Data from two independent experiments were normalized to WT and combined. **b.** CyTOF quantification of proliferating microglia (CD45⁺CD11b⁺CX3CR1⁺P2RY12⁺Ly6C/G⁻CD206⁻CCR2⁻Ki67⁺p16⁻p19⁻p21⁻γH2AX⁻) at day 12 in 7-8-month-old 5xFAD mice treated with isotype control or anti-PD-L1, compared to WT, and in 10–11-month-old mice treated with the same regimens. One-way ANOVA, *P* = 0.003; Fisher’s LSD (uncorrected): WT vs isotype, *P* < 0.001; isotype vs anti-PD-L1, *P* = 0.01. Two independent experiments were normalized to WT, and combined. **c.** CyTOF quantification of homeostatic microglia at day 3 in 7–8-month-old 5xFAD mice treated with isotype control or anti-PD-L1, compared to WT. One-way ANOVA, P = 2.17 × 10⁻⁶; Tukey’s multiple comparisons: WT vs isotype, *P* < 0.0001; WT vs anti-PD-L1, *P* < 0.0001; isotype vs anti-PD-L1, *P* = 0.0434. **d.** CyTOF quantification of proliferating microglia at day 3 in 7–8-month-old 5xFAD mice treated with isotype control or anti-PD-L1, compared to WT. One-way ANOVA, P = 0.0032 ; Fisher’s LSD (uncorrected): WT vs isotype, *P* < 0.001; isotype vs anti-PD-L1, *P* = 0.37 (ns). **e.** FlowJo gating strategy used to define proliferating and senescent microglial populations in the flow cytometry analysis of mouse brains. **f.** Geometric mean expression of PD-L1 in distinct microglial populations from 7-8-month-old 5xFAD brains (n = 3). rMG, resting (homeostatic) microglia; sMG, senescent microglia; pMG, proliferating microglia. One-way ANOVA, *P* = 1.0 × 10⁻⁶. Tukey’s multiple comparisons showed significant differences between rMG and sMG (*P* < 0.0001), rMG and pMG (*P* < 0.0001), and sMG and pMG (*P* = 0.0009).

In contrast, analysis at an earlier timepoint revealed a distinct temporal response. Homeostatic microglia were already increased 3 days after a single injection of anti-PD-L1, though to a lesser extent than observed at 12 days (Fig. 3c), whereas the proportion of proliferating microglia were not affected at this earl time point post-treatment (Fig. 3d). These results suggest that the effect on the proliferating microglia might be secondary to the clearance of the senescent cells. Data obtained at 12 days after treatment were pooled from two independent experiments and normalized to wild-type controls included in each experiment. Data from the 3-daytime-point represent absolute proportions measured within a single experiment.

The differential effect on proliferating versus senescent microglia observed at day 3, prompted us to examine whether these differences may be related to different levels of PD-L1 expressed by these two microglial subsets. We found that both senescent and proliferating microglia express PD-L1, with even slightly higher levels observed in proliferating cells (Fig. 3e, f). We therefore searched for any molecules that might be differentially expressed by senescent microglia and could potentially attract the monocyte-derived macrophages. To this end, we analysed the dataset from Zhou *et al.* (comparing 7-month-old WT and 5XFAD mice)^22^. After quality control (QC) and integration of WT and 5XFAD samples, senescence scores were assigned to all microglia. Despite the relatively young age of these mice (7 months), a small subset of microglia showed a strong senescent transcriptional signature. Notably, approximately 5% of senescent microglia showed marked upregulation of the chemokine Ccl5, a feature that could potentially account for preferential chemotaxis of monocytes toward the senescent cells (Extended Fig. 2).

## Discussion

In the current study, we demonstrate that systemic administration of anti-PD-L1 reduced the burden of senescent microglia in the brain. Furthermore, the treatment reshaped the microglial landscape in 5xFAD mice to resemble that of age-matched WT animals.

Microglia are specialized to detect and respond to deviations from homeostasis. Early in the AD disease process microglia adopt an activated phenotype that supports clearance of misfolded proteins, dying cells, and cellular debris. This activated microglial state includes disease associated microglia (DAM)^23^, which express multiple damage-associated molecule receptors^24^, enabling them to recognize pathogenic patterns, rather than specific proteins^25,26^. In parallel, small subsets of microglia proliferate, potentially enhancing the capacity of the local immune system to contain the pathological conditions^25,26^. Over time, however, microglia progressively lose their clearance functions, and instead become a source of chronic inflammation, in part by acquiring activation states that resemble cellular senescence^27–32^. While many independent studies support a protective role for microglia^33^, their phenotype changes along disease progression ^34^. This temporal and functional heterogeneity emphasizes the limitations of targeting a single microglial protein as a disease-modifying strategy.

Ablation of microglia in an AD mouse model using a selective CSF1R inhibitor was shown to prevent neuronal loss and improve contextual memory^35^. Notably, this intervention did not reduce amyloid-β levels or plaque burden, suggesting that at the disease stage characterized by microglial dysfunction or exhaustion, microglia themselves can act as drivers of cognitive decline. It has been further demonstrated that microglial ablation, followed by their replacement with bone marrow-derived macrophages, may slow or arrest the disease process^35,36^. In that context, our findings that PD-L1 blockade leads to the selective removal of senescent microglia while increasing homeostatic microglial populations, are particularly relevant, as our approach avoids indiscriminate microglial depletion and was previously shown to dampen multiple pathological factors associated with AD, including retarding cognitive loss^13,16–19^.

We further found that the elimination of senescent cells depends upon the CCR2-CCL2 axis, consistent with prior reports that anti-PD-L1 disease modification is mediated through monocyte recruitment to the brain^16,19^. Senescent cells secrete chemokines that attract monocytes to their local niche^39^. Consistent with this, CCL5 levels were previously reported to be markedly elevated in the brains of 5xFAD mice, and treatment with the senolytic compound ABT-737 significantly reduced this chemokine ^13^.

In the present study senescent cell clearance was also observed following treatment with an Fc-null antibody variant currently in clinical development^21^, indicating that this process is independent of classical Fc-mediated effector mechanisms. Recruited monocyte-derived macrophages can directly induce apoptosis of senescent cells via death receptor ligands or oxidative stress, and subsets of these cells may express molecules typically associated with cytotoxic immune linages, such as GZMK and NKG7^42^. In parallel, monocyte-derived macrophages homing to the brain have also been reported to adopt an anti-inflammatory, reparative phenotype^40,41^. Together, these findings support a model in which recruited monocytes contribute to the selective removal, replacement or displacement of senescent microglia through Fc-independent mechanisms rather than direct antibody-mediated cytotoxicity.

In peripheral tissues, senescent cells themselves were shown to express elevated levels of PD-L1 ^43–45^, enabling escape from PD-1-dependent immune surveillance, and thereby accelerating tissue inflammation and aging^43,46^. Consistent with this, blocking the PD-1/PD-L1 pathway effectively eliminates senescent cells in multiple non-CNS tissues^43^. For example, in a lung inflammation model, immune checkpoint blockade removes senescent p16^+^ macrophages by saturating the PD-1 expressed on cytotoxic CD8 T cells^43^.

In the present study, anti-PD-L1 was injected in the periphery, and its levels measured in the cerebrospinal fluid 24 hours after a single intraperitoneal injection. Levels of the Fc-null antibody reached only 0.02-0.05% of serum concentrations (Extended Table 1). These low CNS levels do not support a mechanism that depends on the direct antibody binding within the brain parenchyma, instead supporting an indirect, peripherally-initiated immune mechanism, although additional studies will be required to fully exclude direct local effects.

Taken together, evidence that senescent microglia constitute a major source of brain inflammation, combined with the central role of inflammation as a driver of AD regardless of its primary cause, provides a compelling rationale for anti-PD-L1 therapy as a senolytic-like approach to counter brain aging and age-related dementia. Our findings indicate that the beneficial effects on disease progression result primarily from reduction of inflammation downstream of the elimination of senescent microglia. An Fc-modified anti-PD-L1 antibody is currently being evaluated in a clinical trial in patients with Alzheimer’s disease^21^. Given that PD-1/PD-L1 blockade also reduces senescent cells in multiple peripheral tissues^43^, our results support a model in which peripheral immune system rejuvenation contributes secondary rejuvenation of the brain.

Several limitations of this study should be noted. First, we did not systematically assess non-microglial brain cell populations that may also express PD-L1 or acquire a senescent phenotype in AS, including astrocytes and neurons^47^. Whether these cells contribute to neuroinflammation or are affected by treatment remains to be determined. Finally, because CCR2 is not exclusively expressed by monocytes, contributions from additional CCR2^+^ immune cell populations cannot be excluded.

## Methods

### Animals

Heterozygous 5xFAD mice (C57BL/6J-SJL background) expressing mutant human APP (Swedish K670N/M671L, Florida I716V, London V717I) and mutant PSEN1 (M146L/L286V) under the Thy1 promoter (Tg6799; Jackson Laboratory) were used as an Alzheimer’s disease model. Age-matched non-transgenic littermates served as wild-type controls.

### Antibody adminhistration to mice

Mice were administered anti-PD-L1 (BioXcell), isotype control (BioXcell), or anti-CCR2 by intraperitoneal (IP) injection. Anti-PD-L1 was given as a single dose of 1.5 mg in 400 μl PBS (1×). The isotype control was administered at the same dose and volume. Anti-CCR2 was administered at 400 μg per injection in 200 μl PBS (1×) every 4 days (on days –3, 1, 5 and 9 relative to the anti-PD-L1 or isotype control injection performed on day 0).

### Flow Cytometry

For brain immune-cell analyses, hemispheres were collected after PBS 1x perfusion, cut into small fragments, digested with collagenase (Sigma-Aldrich) and DNase I (Roche), and homogenized to obtain single-cell suspensions. Myelin and debris were removed by centrifugation through a 20% Percoll gradient (Cytiva). Cells were stained with the LIVE/DEAD Fixable Violet Kit, blocked with FcR-blocking Reagent, and incubated with fluorophore-conjugated antibodies (Supplementary Data Table 2).

For intracellular staining, cells were fixed in 4% PFA (Thermo Fisher Scientific) for 10 min at room temperature, washed with FACS buffer, permeabilized with 1× Permeabilization Buffer (Invitrogen) on ice, washed with FACS Buffer, blocked with normal donkey serum, and stained with antibodies against intracellular targets (Supplementary Data Table 1). Data were acquired on a CytoFLEX S flow cytometer at low flow rate.

### Mass Cytometry of Brain (CyTOF)

Brain cells were dissociated, and debris removed as described above. Cells were stained with cisplatin for live/dead discrimination, blocked with FcR-blocking Reagent (BioLegend), and incubated with metal-conjugated surface antibodies (Supplementary Data Table 2). After washing with Maxpar Cell Staining Buffer (Fluidigm), cells were fixed in 4% PFA (Thermo Fisher Scientific), permeabilized with 90% ice-cold methanol, barcoded with palladium-based barcodes, and pooled. Samples were then blocked with normal donkey serum and stained with metal-conjugated intracellular antibodies (Extended Data Table 2). Finally, cells were stained with an iridium intercalator for DNA labelling and acquired on a Helios CyTOF system (Fluidigm).

### Data Analysis

Flow cytometry data were analysed in FlowJo, and CyTOF data were processed in Cytobank and FlowJo. Graphs were prepared, and statistical analyses were performed using GraphPad Prism 10, and schematic illustrations were created with BioRender and Adobe Illustrator.

CyTOF samples were first normalized and de-barcoded using Fluidigm software. Pre-gating and quality control included removal of beads, gating on live cells positive for the ^193Ir/^191Ir DNA intercalator, and exclusion of anomalies based on parameters such as Time, Center/Offset, Width, and Residual (Extended Fig. 1d). For brain samples, CD45⁺ immune cells were gated and subjected to viSNE dimensionality reduction and FlowSOM-based clustering in Cytobank. Because of limited cell numbers and variability in marker expression, CD45⁺-gated brain datasets were also analysed independently in FlowJo.

### Single-cell RNA-seq re-analysis

Published single-cell RNA-seq dataset from Zhou et al. (PMID: 31932797), containing single-cell transcriptomes from 7-month-old WT and 5xFAD mouse brains, was re-processed from raw matrices using Seurat v5 (Satija Lab). Data were filtered following quality thresholds consistent with the original publication and integrated across genotypes using standard Seurat workflows. Microglia subsets were identified based on published annotations. A senescence signature score was computed using canonical senescence-associated genes (including *Cdkn1a*, *Cdkn2a* and *Gadd45g*), and microglia within the top-scoring fraction were classified as high-senescence. Differential expression between high-senescence and other microglia was performed using SCTransform-normalised values and *FindMarkers* (adjusted *P* < 0.05).

---

**Extended Data Fig. 1.**
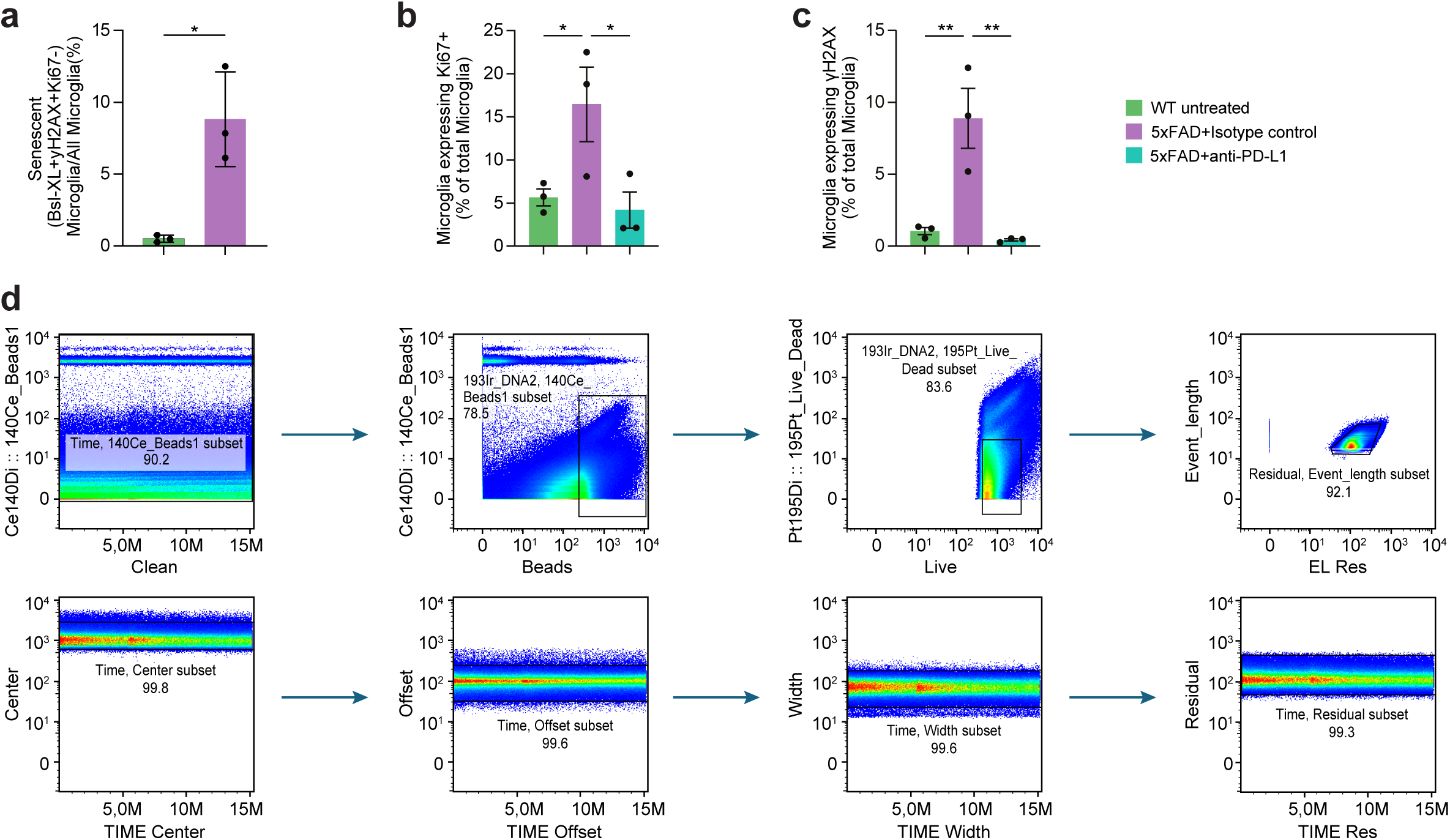
Flow cytometry and CyTOF gating strategies for microglial analysis. **a.** Flow cytometry quantification of senescent microglia (CD45⁺CD11b⁺Ly6C⁻γH2AX⁺BSL-xl⁺Ki67⁻) in 8–9-month-old 5xFAD mice compared to WT controls. Two-tailed t-test, *P* = 0.0478. **b.** Flow cytometry quantification of proliferating microglia (CD45⁺CD11b⁺Ly6C⁻Ki67⁺) at day 14 in 8–9-month-old 5xFAD mice treated with anti-PD-L1 or isotype control, compared to WT. One-way ANOVA, *P* = 0.04; Fisher’s LSD (uncorrected): isotype vs WT, *P* = 0.04; isotype vs anti-PD-L1, *P* = 0.02. **c.** Flow cytometry quantification of γH2AX⁺ microglia at day 14 in 8–9-month-old 5xFAD mice treated with anti-PD-L1 or isotype control, compared to WT. One-way ANOVA, *P* = 0.005; Fisher’s LSD (uncorrected): isotype vs WT, *P* = 0.004; isotype vs anti-PD-L1, *P* = 0.003. **d.** CyTOF pre-gating workflow for initial data clean-up in FlowJo.

**Extended Data Fig. 2.**
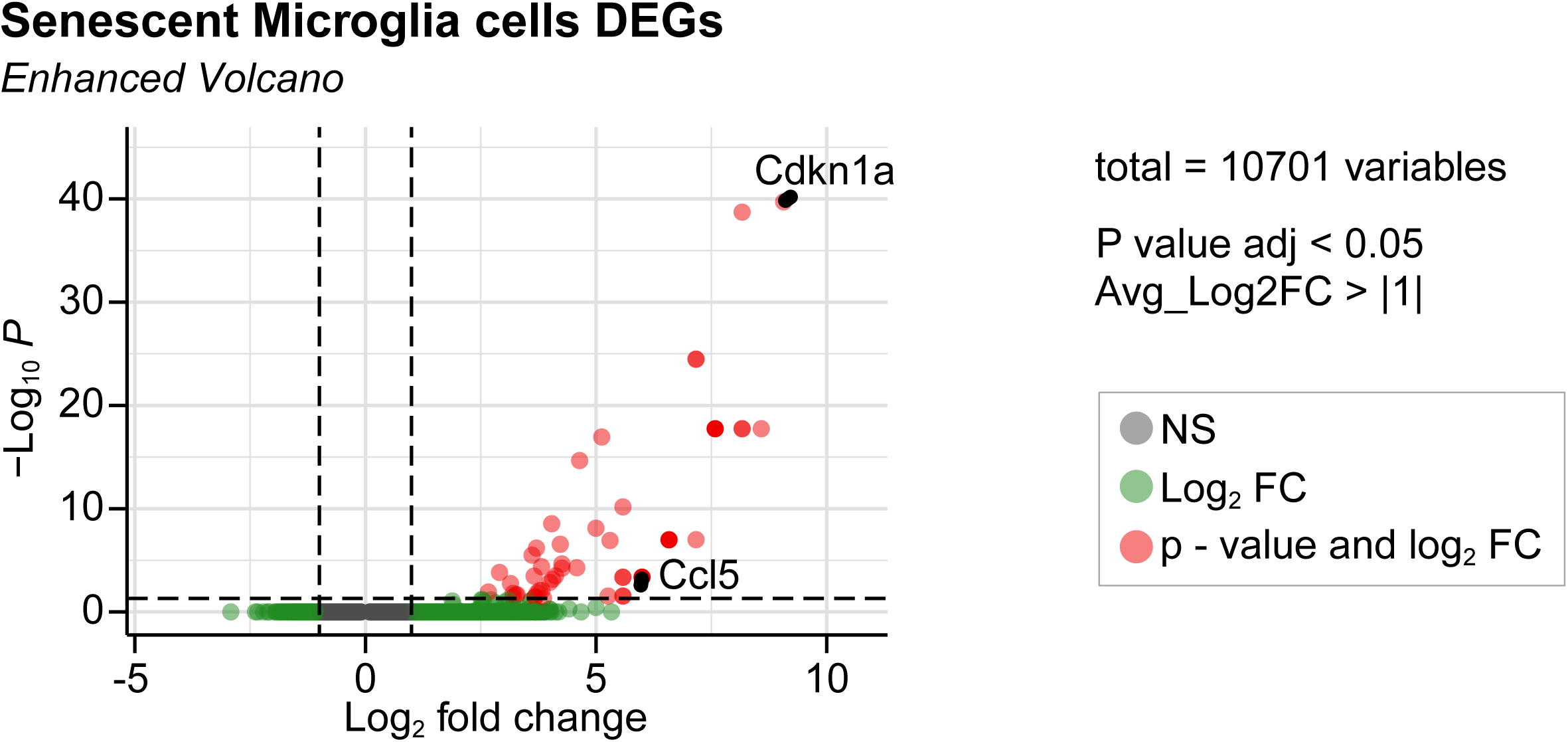
scRNA-seq analysis of mouse microglia. Volcano plot showing significant upregulation (adj. *P* < 0.05) of CCL5 in microglia classified as “high-senescence,” defined by elevated expression of core senescence-associated genes (e.g. *Cdkn1a*).

## Extended Data

**Table 1.** Serum and CSF concentration of anti-PD-L1 Fc-null antibody, 24 hours after single injection. LLOQ: 0000137ug/ml.

| Mouse | Injected dose<br>(i.p.) | Time | CSF concentration<br>(µg/ml) | Serum concentration<br>(µg/ml) | %CSF/Serum |
| --- | --- | --- | --- | --- | --- |
| 1 | 1.5 mg/mouse | 24 hr | 0.033244 | 103.9082 | 0.032% |
| 2 | 1.5 mg/mouse | 24 hr | 0.046673 | 85.99364 | 0.054% |
| 3 | 0.1 mg/mouse | 24 hr | 0.001144 | 5.325588 | 0.021% |
| 4 | 0.1 mg/mouse | 24 hr | <0.0000137 | 0.029242 | <0.047% |

## Acknowledgments

This study was supported by Thompson foundation a grant from the ERNA-NET (given to MS).

## Contributions

Y.A. and M.S. conceptualized the study. Y.A. designed and performed the CyTOF and flow cytometry experiments. M.A., B.N., H.P., A.K. and A.I. contributed to experimental execution and study design. M.S. and Y.A. wrote the manuscript. V.K. contributed to project conceptualization. H.A. performed the analysis of the sequencing datasets. T.M.S. assisted with CyTOF panel design and the technical aspects of mass cytometry.

## References

1. Castellani, G., Croese, T., Peralta Ramos, J. M. & Schwartz, M. Transforming the understanding of brain immunity. Science 380, eabo7649 (2023).

2. Basurco, L., Abellanas, M. A., Purnapatre, M., Antonello, P. & Schwartz, M. Chronological versus immunological aging: Immune rejuvenation to arrest cognitive decline. Neuron 113, 140–153 (2025).

3. Cantone, A. F. et al. Rebalancing Immune Interactions within the Brain-Spleen Axis Mitigates Neuroinflammation in an Aging Mouse Model of Alzheimer’s Disease. J Neuroimmune Pharmacol 20, 15 (2025).

4. Oh, H. S.-H. et al. Plasma proteomics links brain and immune system aging with healthspan and longevity. Nat Med 31, 2703–2711 (2025).

5. Wyss-Coray, T. Inflammation in Alzheimer disease: driving force, bystander or beneficial response? Nat Med 12, 1005–15 (2006).

6. Zhang, Q. et al. Neuroinflammation in Alzheimer’s disease: insights from peripheral immune cells. Immun Ageing 21, 38 (2024).

7. Zilka, N. et al. Who fans the flames of Alzheimer’s disease brains? Misfolded tau on the crossroad of neurodegenerative and inflammatory pathways. J Neuroinflammation 9, 47 (2012).

8. Selkoe, D. J. Treatments for Alzheimer’s disease emerge. Science 373, 624–626 (2021).

9. Hansen, D. V, Hanson, J. E. & Sheng, M. Microglia in Alzheimer’s disease. J Cell Biol 217, 459–472 (2018).

10. Hu, Y. et al. Replicative senescence dictates the emergence of disease-associated microglia and contributes to Aβ pathology. Cell Rep 35, 109228 (2021).

11. Mishra, R. Cellular Senescence in the Aging Brain. in Cellular Senescence and Brain Aging 55–78 (Springer Nature Singapore, Singapore, 2025). doi:10.1007/978-981-96-8873-9_4.

12. Onyango, I. G., Jauregui, G. V, Čarná, M., Bennett, J. P. & Stokin, G. B. Neuroinflammation in Alzheimer’s Disease. Biomedicines 9, (2021).

13. Rachmian, N. et al. Identification of senescent, TREM2-expressing microglia in aging and Alzheimer’s disease model mouse brain. Nat Neurosci 27, 1116–1124 (2024).

14. Gross, P. S. et al. Senescent-like microglia limit remyelination through the senescence associated secretory phenotype. Nat Commun 16, 2283 (2025).

15. Li, W., Yong-Yan, X., Jia-Xin, M., Shu-Chao, G. & Li-Ping, H. Senescent microglia: The hidden culprits accelerating Alzheimer’s disease. Brain Res 1851, 149480 (2025).

16. Rosenzweig, N. et al. PD-1/PD-L1 checkpoint blockade harnesses monocyte-derived macrophages to combat cognitive impairment in a tauopathy mouse model. Nat Commun 10, 465 (2019).

17. Ben-Yehuda, H. et al. Key role of the CCR2-CCL2 axis in disease modification in a mouse model of tauopathy. Mol Neurodegener 16, 39 (2021).

18. Dvir-Szternfeld, R. et al. Alzheimer’s disease modification mediated by bone marrow-derived macrophages via a TREM2-independent pathway in mouse model of amyloidosis. Nat Aging 2, 60–73 (2022).

19. Baruch, K. et al. PD-1 immune checkpoint blockade reduces pathology and improves memory in mouse models of Alzheimer’s disease. Nat Med 22, 135–7 (2016).

20. Xing, Z. et al. Influenza vaccine combined with moderate-dose PD1 blockade reduces amyloid-β accumulation and improves cognition in APP/PS1 mice. Brain Behav Immun 91, 128–141 (2021).

21. Baruch, K. et al. IBC-Ab002, an anti-PD-L1 monoclonal antibody tailored for treating Alzheimer’s disease. Alzheimer’s & Dementia 16, (2020).

22. Zhou, Y. et al. Human and mouse single-nucleus transcriptomics reveal TREM2-dependent and TREM2-independent cellular responses in Alzheimer’s disease. Nat Med 26, 131–142 (2020).

23. Huang, Y. et al. Microglia use TAM receptors to detect and engulf amyloid β plaques. Nat Immunol 22, 586–594 (2021).

24. Rolls, A. et al. Toll-like receptors modulate adult hippocampal neurogenesis. Nat Cell Biol 9, 1081–8 (2007).

25. Baligács, N. et al. Homeostatic microglia initially seed and activated microglia later reshape amyloid plaques in Alzheimer’s Disease. Nat Commun 15, 10634 (2024).

26. He, C., Chen, B., Yang, H. & Zhou, X. The dual role of microglia in Alzheimer’s disease: from immune regulation to pathological progression. Front Aging Neurosci 17, (2025).

27. Krasemann, S. et al. The TREM2-APOE Pathway Drives the Transcriptional Phenotype of Dysfunctional Microglia in Neurodegenerative Diseases. Immunity 47, 566–581.e9 (2017).

28. Habib, N. et al. Disease-associated astrocytes in Alzheimer’s disease and aging. Nat Neurosci 23, 701–706 (2020).

29. Udeochu, J. C. et al. Tau activation of microglial cGAS-IFN reduces MEF2C-mediated cognitive resilience. Nat Neurosci 26, 737–750 (2023).

30. Xie, X. et al. Activation of innate immune cGAS-STING pathway contributes to Alzheimer’s pathogenesis in 5×FAD mice. Nat Aging 3, 202–212 (2023).

31. Carling, G. K. et al. Alzheimer’s disease-linked risk alleles elevate microglial cGAS-associated senescence and neurodegeneration in a tauopathy model. Neuron 112, 3877–3896.e8 (2024).

32. Deczkowska, A. et al. Mef2C restrains microglial inflammatory response and is lost in brain ageing in an IFN-I-dependent manner. Nat Commun 8, 717 (2017).

33. Yin, Z. et al. Identification of a protective microglial state mediated by miR-155 and interferon-γ signaling in a mouse model of Alzheimer’s disease. Nat Neurosci 26, 1196–1207 (2023).

34. Pilat, D. J. et al. The gain-of-function TREM2-T96K mutation increases risk for Alzheimer’s disease by impairing microglial function. Neuron 10.1016/j.neuron.2025.09.032 (2025) doi:10.1016/j.neuron.2025.09.032.

35. Wu, J. et al. Microglia replacement halts the progression of microgliopathy in mice and humans. Science (1979) 389, (2025).

36. Zhou, K., Wang, Y., Xu, Y., Zhang, Y. & Zhu, C. Exploring microglial replacement: From disease models to clinical translation. Brain Behav Immun 129, 179–185 (2025).

37. Abellanas, M. A., Purnapatre, M., Burgaletto, C. & Schwartz, M. Monocyte-derived macrophages act as reinforcements when microglia fall short in Alzheimer’s disease. Nat Neurosci 28, 436–445 (2025).

38. Behmoaras, J. & Gil, J. Similarities and interplay between senescent cells and macrophages. J Cell Biol 220, (2021).

39. Han, X. et al. Potential Regulators of the Senescence-Associated Secretory Phenotype During Senescence and Aging. J Gerontol A Biol Sci Med Sci 77, 2207–2218 (2022).

40. Han, X. et al. Potential Regulators of the Senescence-Associated Secretory Phenotype During Senescence and Aging. J Gerontol A Biol Sci Med Sci 77, 2207–2218 (2022).

41. Elder, S. S. & Emmerson, E. Senescent cells and macrophages: key players for regeneration? Open Biol 10, 200309 (2020).

42. Turiello, R. et al. NKG7 is a Stable Marker of Cytotoxicity Across Immune Contexts and Within the Tumor Microenvironment. Eur J Immunol 55, e51885 (2025).

43. Wang, T.-W. et al. Blocking PD-L1-PD-1 improves senescence surveillance and ageing phenotypes. Nature 611, 358–364 (2022).

44. Pippin, J. W. et al. Upregulated PD-1 signaling antagonizes glomerular health in aged kidneys and disease. Journal of Clinical Investigation 132, (2022).

45. Onorati, A. et al. Upregulation of PD-L1 in Senescence and Aging. Mol Cell Biol 42, (2022).

46. Majewska, J. et al. p16-dependent increase of PD-L1 stability regulates immunosurveillance of senescent cells. Nat Cell Biol 26, 1336–1345 (2024).

47. Zhu, J., Wu, C. & Yang, L. Cellular senescence in Alzheimer’s disease: from physiology to pathology. Transl Neurodegener 13, 55 (2024).

