## Supplemental Table 1 and 2 for "Systemic administration of PD-L1 blocking antibodies leads to removal of senescent microglia"

**Supplementary**

**Table 1**

Flow Cytometry antibodies

| N | AB | Fluorochrome | Clone |
|---|---|---|---|
| 1 | Ki67 | BV650 | 11F6, BioLegend |
| 2 | BSL-xL | PE-Cy7 | 54H6, Cell Signalling |
| 3 | yH2AX | PE | N1-431, BD Pharmingen |
| 4 | CD11b | APC | M1/70, BioLegend |
| 5 | CD45 | BV421 | 30-F11, Biolegend |
| 6 | L/D | BV510 | Aqua fluorescent reactive dye, Invitrogen |
| 7 | Ly6c | PE-Dazzle | HK1.4, Biolegend |
| 8 | PD-L1 | PE-Cy5 | 10F.9G2, Biolegend |

**Table 2**

CyTOF antibodies

| N | Ab | Clone | Metal |
|---|---|---|---|
| 1 | CD11b | M1/70 | 143Nd |
| 2 | CD206 | C068C2 | 169Tm |
| 3 | CD44 | IM7 | 150Nd |
| 4 | P2ry12 | SXM30 | 160Yb |
| 5 | CD11c | N418 | 209Bi |
| 6 | Ly6G/C | RB6-8C5 | 141Pr |
| 7 | Cx3cr1 | SA011F11 | 164Dy |
| 8 | CD38 | 90 | 175Lu |
| 9 | CD45 | 30-F11 | 89Y |
| 10 | CCR2 | 8C3.1 | 171Yb |
| 11 | TREM2 | 237920 | 153Eu |
| 12 | CD9 | kmc8 | 158Gd |
| 13 | CD39 | 24dms1 | 142Nd |
| 14 | γH2AX | NB100-384 | 149Sm |
| 15 | P21 | SXM30 | 152Sm |
| 16 | P19 | EPR20418 | 166Er |
| 17 | P16 | 2D9A12 | 167Er |
| 18 | Ki-67 | B56 | 168Er |
